# Alzheimer’s disease-like features in resting state EEG/fMRI of cognitively intact and healthy middle-aged *APOE*/*PICALM* risk carriers

**DOI:** 10.1101/2024.06.20.599857

**Authors:** Patrycja Dzianok, Jakub Wojciechowski, Tomasz Wolak, Ewa Kublik

## Abstract

**Introduction:** Genetic susceptibility is a primary factor contributing to etiology of late-onset Alzheimer’s disease (LOAD). The exact mechanisms and timeline through which *APOE*/*PICALM* influence brain functions and contribute to LOAD remain unidentified. This includes their effects on individuals prior to the development of the disease.

**Methods:** *APOE*/*PICALM* alleles were assessed to determine the genetic risk of LOAD in 79 healthy, middle-aged participants who underwent EEG and fMRI recordings. The resting-state signal was analyzed to estimate relative spectral power, complexity (Higuchi’s algorithm), and connectivity (coherence in EEG and ICA-based connectivity in fMRI).

**Results:** The main findings indicated that individuals at risk for LOAD exhibited reduced signal complexity and the so-called “slowing of EEG” which are well-known EEG markers of AD. Additionally, these individuals showed altered functional connectivity in fMRI (within attention related areas).

**Discussion:** Risk alleles of *APOE/PICALM* may affect brain integrity and function prior to the onset of the disease

## Background

Understanding the early development of LOAD is crucial for its effective diagnosis, prevention, and treatment. The apolipoprotein E gene (*APOE*) is widely recognized as the predominant genetic factor influencing LOAD. It has three isoforms: ε4, ε3, and ε2. The ε4 increases the risk of developing AD by 4-12 times compared to non-carriers [1,2]. In contrast, the ε3 isoform appears to have no effect on disease risk, while the ε2 isoform is occasionally associated with a reduced risk [3]. Patients with the homozygous ε4 allele typically exhibit an earlier onset of disease [4], a phenomenon that is also observed in familial forms of early-onset Alzheimer’s [5]. Genome-wide association studies (GWAS) have identified numerous additional risk genes for Alzheimer’s disease (AD), including 42 new loci reported in a 2022 study [6]. Among these, the gene encoding phosphatidylinositol binding clathrin assembly protein (*PICALM*) has been repeatedly identified as a significant risk factor for AD [6–8]. The *PICALM* G allele is more prevalent among AD patients, whereas the A allele is thought to either decrease the risk of AD or have no effect. Furthermore, potential interactions between the *APOE* and *PICALM* genes have been observed [7]. Some studies suggest that the combined presence of these genes influence brain atrophy and diminishes cognitive performance in early AD patients [9]. Both genes are also implicated in amyloid pathology, a common pathway in the development of AD [10]. They have never been studied together in a non-demented population.

EEG is one of the most promising tools in search for LOAD diagnostic markers [11,12], as it has high availability, low cost and non-invasiveness. The most common protocol used in AD patients is the “resting-state” protocol, as it is brief and does not require participants to engage in any specific task. Most studies use eyes-closed condition [13–18], as open eyes resting-state is often characterized by a EEG desynchronization in common bands of interests. A number of changes in spontaneous EEG has been shown in patients with AD, other dementias and mild cognitive impairment (MCI). The most recognized AD hallmark measured with EEG is the so-called “slowing of EEG” [16,17,19–21], i.e. increased amplitude/power of slow waves as delta (∼1-3 Hz) [13,15–17] and theta (∼4-7 Hz) [13,14,16,17] and decreased amplitude/power of alpha band (∼8-12 Hz) [13,15–17]. Higher frequency (beta, gamma) is rarely reported to be changed across AD continuum or has no effect [13–17]. Signal complexity is another frequently used EEG measure due to the complex and nonlinear dynamics of brain signals. MCI/AD patients have lower signal complexity than healthy controls [20,22,23]. Resting-state protocol allows also for studying functional connectivity (FC). This can be done using EEG signal, but more robust way is to use functional magnetic resonance imaging (fMRI). LOAD tends to be associated with a reduction in functional connectivity in posterior DMN [24–26]. This region was shown to be involved in many actions like memory, introspection, mind-wandering, the generation of spontaneous thought, the maintenance of the sense of self, and the integration of information across different cognitive domains [27]. In fMRI studies, functional connectivity is typically measured using either seed-based or ICA approaches. ICA was used extensively in rs-fMRI studies on AD or individuals at risk with different genetic burden [26,28–32]. As the LOAD etiology is multifaced, it is also important to take into account the neuropsychological, health related and lifestyle aspect. AD patients are characterized by increased apathy/depression, impaired emotional control, or personality changes [33] and other lifestyle and health factors are linked to the greater dementia risk [34].

Studies indicate that adjusting lifestyle and beginning interventions early in the disease process can alter (in some cases) the disease’s progression [11,35,36]. Additionally, modern clinical trials targeting potentially disease-modifying medications focus on the prodromal phases of AD. Thus, understanding the risk-genes influence on health and brain is a key challenge in AD research.

## Material and methods

### Participants and genetic screening

We tested 79 non-demented middle-aged adults during the neuroimaging phase of our study, which was part of a larger research project [37]. EEG session was conducted in the EEG laboratory at the Nencki Institute of Experimental Biology PAS (Poland) and MRI/fMRI session in the Bioimaging Research Center, Institute of Physiology and Pathology of Hearing (Poland). A larger cohort (N = 200) underwent genetic screening and completed questionnaires on demographics, health, and psychometric assessments. From this cohort, 79 subjects were selected based on their genetic scores to form the experimental groups. Exclusion and inclusion criteria were already described in the data note article regarding our database [37]. AD risk genes, the *APOE* (rs429358/rs7412) and *PICALM* (rs3851179) alleles were determined using the traditional Sanger sequencing protocol, which was outsourced to the certified third party company. The participants in the genetic-based research groups were matched based on age, gender, education, and various health factors, particularly those influencing dementia risk. The groups were constructed based on *APOE*/*PICALM* risk: *APOE*-ε4/*PICALM* GG non-carriers (referred to as “N”), single-risk carriers (*APOE*-ε4 carriers without the *PICALM* risky GG alleles, referred to as “A+P-“), and double-risk carriers (*APOE*-ε4 carriers with the *PICALM* risky GG alleles, referred to as “A+P+”).

Several participants withdrew from the study for reasons such as MRI contraindications, and some data were lost due to technical issues. The exact number of participants for each experiment is as follows:

— Health and psychometric tests were completed by all 79 participants (details of missing data in various questionnaires are provided in the Results section Tab. 1).
— EEG data: N = 78 participants (N group: 31, A+P-group: 27, A+P+ group: 20).
— MRI/fMRI data: N = 69 participants (N group: 27, A+P-group: 24, A+P+group: 18).

**Table 1.** Demographic characteristics, health and psychometric tests assessment: descriptive statistics and group differences.

|  | N | A+P- | A+P+ | *p*-value |
| --- | --- | --- | --- | --- |
| **Demographic and health measures** |  |  |  |  |
| Age [years; M±SD] | 54.77 ± 2.92 | 55.70 ± 3.27 | 55.62 ± 3.31 | .47 & |
| Gender [F/M] | 15/16 | 14/13 | 10/11 | .95 ^ |
| Education: secondary [n] | 3 (10.71%) | 4 (17.39%) | 1 (5%) | .45 ^ |
| Education: partial higher [n] | 2 (7.14%) | 2 (8.70%) | 0 (0%) |  |
| Education: higher [n] | 23 (82.14%) | 17 (73.91%) | 19 (95%) |  |
| Handedness EHI [M±SD] | 87.85 ± 20.95 | 86.67 ± 18.81 | 90.16 ± 6.75 | .67 & |
| AUDIT scores [M±SD] | 3.76 ± 3.52 | 4.19 ± 2.91 | 3.87 ± 2.09 | .61 & |
| Smokers [n] | 2 (6.45%) | 5 (18.52%) | 2 (9.52%) | .24 ^ |
| Former smokers [n] | 7 (22.58%) | 10 (37.04%) | 4 (19.05%) |  |
| Diabetes [n] | 0 (0%) | 0 (0%) | 1 (4.76%) | .25 ^ |
| Hypertension [n] | 6 (19.36%) | 9 (33.33%) | 5 (23.81%) | .47 ^ |
| Thyroid [n] | 2 (6.67%) | 3 (11.11%) | 2 (9.52%) | .84 ^ |
| Other diseases [n] | 7 (23.33%) | 6 (22.22%) | 5 (23.81%) | .99 ^ |
| Allergies [n] | 7 (25.00%) | 5 (19.23%) | 4 (21.05%) | .87 ^ |
| BMI [M±SD] | 26.70 ± 4.35 | 27.24 ± 5.48 | 28.58 ± 5.55 | .47 & |
| Drugs [n] | 19 (61.29%) | 17 (62.96%) | 8 (38.10%) | .17 ^ |
| NSAID [n] | 7 (22.58%) | 11 (40.74%) | 5 (23.81%) | .26 ^ |
| Learning difficulties [n] | 4 (13.33%) | 3 (12.50%) | 1 (5.26%) | .65 ^ |
| Family history of dementia [n] | 8 (25.81%) | 14 (51.85%) | 8 (38.10%) | .19 ^ |
| **Psychometric tests assessment [M±SD]** |  |  |  |  |
| BDI | 4.61 ± 4.90 | 8.70 ± 6.51 | 6.62 ± 6.98 | < .05 & |
| SES | 33.19 ± 3.99 | 30.59 ± 4.05 | 32.29 ± 3.76 | < .05 # |
| Mini-Cope-1 | 2.26 ± 0.33 | 2.06 ± 0.48 | 2.21 ± 0.49 | .18 & |
| Mini-Cope-2 | 0.60 ± 0.43 | 0.85 ± 0.40 | 0.74 ± 0.40 | .07 # |
| Mini-Cope-3 | 1.74 ± 0.56 | 1.60 ± 0.64 | 1.51 ± 0.71 | .48 & |
| Mini-Cope-4 | 1.02 ± 0.39 | 1.24 ± 0.33 | 1.07 ± 0.43 | .11 & |
| Mini-Cope-5 | 0.52 ± 0.82 | 0.32 ± 0.72 | 1.26 ± 1.19 | .16 # |
| Mini-Cope-6 | 2.14 ± 0.48 | 2.09 ± 0.65 | 2.07 ± 0.73 | .93 |
| Mini-Cope-7 | 0.86 ± 0.63 | 0.94 ± 0.70 | 1.10 ± 0.65 | .35 & |
| **NEO-NEU** | **14.23 ± 5.91** | **19.96 ± 9.44** | **15.62 ± 8.18** | **< .05 &** |
| NEO-EXT | 27.90 ± 6.65 | 25.48 ± 7.06 | 26.05 ± 6.85 | .38 # |
| NEO-OPE | 30.77 ± 5.31 | 30.82 ± 6.04 | 31.43 ± 6.08 | .91 # |
| NEO-AGR | 33.58 ± 5.83 | 30.96 ± 6.60 | 32.81 ± 6.17 | .27 # |
| NEO-CON | 32.68 ± 6.86 | 29.33 ± 6.29 | 30.86 ± 7.65 | .19 # |
| RAVEN | 53.87 ± 3.19 | 52.11 ± 6.04 | 53.05 ± 4.08 | .73 & |
| CVLT-1 | 61.29 ± 8.86 | 63.63 ± 8.13 | 59.76 ± 8.89 | .39 & |
| CVLT-2 | 9.39 ± 1.69 | 9.70 ± 2.02 | 8.91 ± 1.90 | .41 & |
| CVLT-3 | 8.38 ± 1.91 | 8.41 ± 2.31 | 7.81 ± 1.85 | .36 & |
| CVLT-4 | 12.97 ± 2.65 | 12.74 ± 2.49 | 12.86 ± 3.10 | .78 & |
| CVLT-5 | 13.42 ± 1.96 | 13.59 ± 1.65 | 12.38 ± 2.27 | .99 & |
| CVLT-6 | 13.68 ± 2.36 | 13.33 ± 2.69 | 13.24 ± 3.24 | .95 & |
| CVLT-7 | 13.71 ± 2.02 | 13.67 ± 2.00 | 13.38 ± 2.96 | .98 & |
| CVLT-8 | 2.94 ± 3.85 | 4.63 ± 5.51 | 4.67 ± 4.80 | .18 & |
| CVLT-9 | 1.39 ± 1.94 | 0.85 ± 1.35 | 1.14 ± 1.49 | .33 & |
| CVLT-10 | 1.19 ± 1.87 | 0.78 ± 1.22 | 1.43 ± 3.08 | .77 & |
| CVLT-11 | 15.32 ± 1.30 | 15.19 ± 1.24 | 15.45 ± 0.89 | .63 & |
| CVLT-12 | 0.52 ± 1.09 | 0.44 ± 1.01 | 1.05 ± 2.09 | .41 & |

The study was approved by the local bioethics committee (Bioethics Committee of the Nicolaus Copernicus University in Toruń functioning at Collegium Medicum in Bydgoszcz, Poland). Written informed consent was provided by all participants and all participants received cash remuneration.

### Demographics, health, and psychometric assessment

Participants provided standard demographic information along with details about their health status (e.g., diabetes, hypertension, etc.; all measures are presented in Tab. 1 in Results section). They were then assessed using a comprehensive battery of psychometric tests to evaluate basic characteristics linked to increased dementia risk or typically found in dementia patients. These tests included:

— Depression/apathy (measured by Beck’s Depression Inventory, BDI)
— Self-esteem (measured by Rosenberg’s Self-Esteem Scale, SES)
— Stress and stress coping strategies (measured by Mini-Cope Questionnaire)
— Personality (measured by NEO-FFI Personality Inventory)
— Intelligence (measured by Raven’s Progressive Matrices – standard/classic version, RPM)
— Memory (measured by California Verbal Learning Test, CVLT)

Additionally, alcohol use was measured by the Alcohol Use Disorders Identification Test (AUDIT; a threshold of ≥ 8 points could indicate unhealthy alcohol usage) and handedness by the Edinburgh Handedness Inventory (EHI).

### Data acquisition

Both EEG and fMRI experiments included an eyes-closed resting-state condition. To ensure participants were rested, EEG sessions were conducted exclusively in the morning and early afternoon. The sessions took place in a comfortable room with dim lighting, where participants sat in a comfortable chair with armrests and faced a monitor. A researcher supervised the study remotely via computer and online LAN camera. Participants were instructed to relax, avoid thinking about anything specific, and remain still. EEG was recorded for 6 minutes using the extended 10-20 international system for electrode placement (Fig. 1), with 128 active electrodes (actiCAP, Brain Products, Munich, Germany) on a Brain Products EEG system. The online reference was set to FCz. At the end of the session, a handheld CapTrak 3D scanner (Brain Products) was used to obtain accurate electrode locations. Impedance was kept as low as possible (average 7.84±3.15 kΩ) through skin rubbing and gel application (Supervisc, extra viscous gel). A low-pass filter was set to 280 Hz, and no high-pass or Notch filters were used during recording. The sampling rate was 1000 Hz.

**Figure 1.**
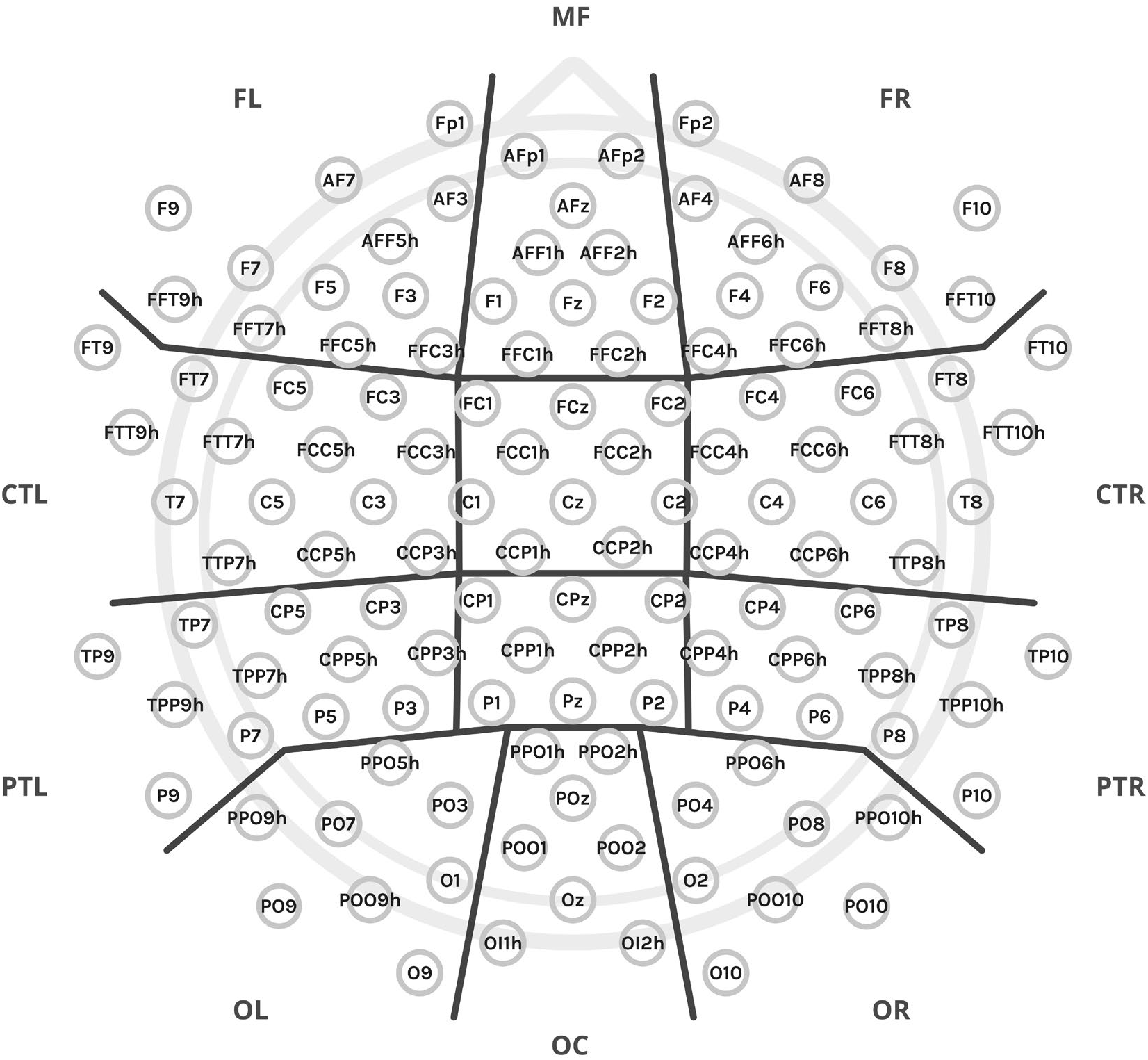
The experiment employed a setup of 128 electrodes, which were organized into anatomical clusters for data analysis, as denoted by the thick black lines: MF – midfrontal, FL – frontal left, FR – frontal right, C – central, CTL – central-temporal left, CTR – central-temporal right, PR – parietal central, PTL – parietal-temporal left, PTR – parietal-temporal right, OC – occipital central, OL – occipital left, and OR – occipital right. Additionally, two midline clusters, not marked in the figure, include C – central (comprising FCz, Cz, and neighboring electrodes) and CP – central-parietal (comprising CPz, Pz, and neighboring electrodes).

MRI/fMRI experiments were performed on a 3T Siemens Prisma FIT scanner (Siemens Medical Systems, Erlangen, Germany) equipped with a 64-channel phased-array RF head coil. The acquisition parameters were as follows: multi-band (slice acceleration factor = 8) EPI sequence, repetition time (TR) = 0.8 s, echo time (TE) = 0.038 s, slice thickness = 2 mm, 72 slices, IPAT = 1, FOV = 216×216 mm, 52° flip angle, voxel size = 2×2×2 mm, and acquisition time (TA) = 7:30. Each subject underwent two resting-state sequences, one with Anterior-Posterior encoding phase and the other with Posterior-Anterior encoding phase. Structural T1-weighted 3D MP-Rage images were acquired with the following parameters: TR = 2400 ms, TI = 1000 ms, TE = 2.74 ms, 8° flip angle, FOV = 256×256 mm, voxel size = 0.8×0.8×0.8 mm, and TA = 6:52 minutes.

### Data preprocessing

The EEG data were preprocessed using the EEGLAB toolbox [38] within MATLAB 2022a. For each participant, standard electrode positions in EEG data files were replaced with individual positions obtained from the CapTrak localizer. The data were downsampled to 250 Hz and filtered within the range of 0.1-40 Hz using standard filter parameters from the toolbox. Additional files were saved with filtering specifically set to 1-40 Hz for subsequent ICA. Channels with excessive noise were removed based on the EEGLAB clean raw data algorithm, which utilizes criteria such as no-signal/flat line, channel correlation, and line noise, as well as through visual inspection. On average, 5.33 channels out of 127 were removed per participant, and the removed channels were interpolated. An average reference was applied, and the initial reference electrode (FCz) was restored and included in the data. The data were segmented into non-overlapping epochs, and those containing excessive artifacts were removed using the ASR algorithm (artifact subspace reconstruction bad burst correction) and further visual inspection. On average, 2.68 epochs per participant were removed. ICA was then applied to detect and separate components with evident physiological artifacts (e.g., eye-blink, muscle, ECG artifacts), resulting in the removal of an average of 5.81 components per participant. The data were visually inspected again, and if necessary, additional cleaning of epochs was performed, with an average of 1.65 additional epochs removed per participant.

Preprocessing of the MRI/fMRI data was performed using SPM12 (Wellcome Trust Centre for Neuroimaging, London, UK) and FSL. Functional data were first realigned, followed by correction of spatial distortion from the encoding phase using FSL’s topup function. The structural T1-weighted image was co-registered with the functional images, segmented, and normalized to the common 1-mm isometric MNI space. Transformation parameters obtained from this process were then applied to the functional images after resampling to a 2-mm isometric voxel size. A 6 mm Gaussian kernel (full width half maximum, FWHM) was used for spatial smoothing. Functional data were filtered in the 0.008 to 0.09 Hz band range and denoised using ArtToolbox, as implemented in CONN [39], with ‘intermediate settings’ (Global-signal z-value < 5; motion < 0.9 mm). Additionally, the COMPCOR [40] approach was utilized on white matter and cerebrospinal fluid signals to generate nuisance regressors related to physiological artifacts (6 PCA components for each mask).

### Data analysis

The analysis of the EEG data was performed using MATLAB. For each participant, Welch’s power spectral density estimate was computed at each channel with a 4-second window and 50% overlap (*pwelch* MATLAB function; spectral resolution 0.25 Hz). The average power for the delta (0.5-4 Hz), theta (4-7 Hz), alpha-1 (7.5-9.5 Hz), and alpha-2 (10-12 Hz) frequency bands was then calculated using the *bandpower* MATLAB function. Relative average band power was calculated by dividing the band scores by the total power of the signal in the 1-30 Hz range. Global relative band powers (average of all 128 electrodes) and regional relative band powers (average of selected electrodes) were computed. For regional power estimation, electrodes were grouped into 12 anatomical regions of interest (ROIs)/clusters: midfrontal (MF), left and right frontal (FL and FR), left and right central-temporal (CTL and CTR), left and right parietal-temporal (PTL and PTR), and left and right occipital (OL and OR) (Fig. 1). To ensure an approximately normal distribution of data for statistical analysis, the results were logit-transformed using the function *t*(*x*) = *log*(*x*(1 − *x*)) [41].

The EEG signal complexity was calculated using Higuchi’s fractal dimension (HFD) algorithm [42,43], which is renowned for its accurate estimation of signal fractal dimension in electrophysiology data and for its computational efficiency [44]. Among various algorithms, HFD consistently provides the most precise fractal dimension estimations [45,46] and has been effective in distinguishing between Alzheimer’s disease patients and healthy subjects [23,47]. The fractal dimension ranges from 1 to 2, with higher values indicating more complex signals. This value depends on the algorithm’s tuning parameter, *k*_max_, which can be defined using several approaches. We selected *k*_max_ using the plateau criterion, which has proven efficient for EEG data, though it is not always recommended for other data types [48]. This approach, validated for various data types [48,49], identifies the best discrimination between predefined groups. To determine the appropriate *k*_max_ for calculating group difference statistics, we first computed the absolute percentage change between average HFD values at consecutive *k* values to detect the plateau. A threshold of 0.1% was used to find the start of the function minima. The first minimum was at *k*_max_ = 36, but subsequent values exceeded the threshold. The second minimum, at *k*_max_ = 62, marked the start of the function plateau, where percentage changes remained below 0.1%. Next, we calculated the distance metric (pairwise difference) between the three groups over the plateau values (from *k*_max_ = 62 to *k*_max_ = 100). The sum of differences for each *k*_max_ was then computed. The *k*_max_ = 82 corresponded to the largest sum of differences among the groups, indicating the point where the groups were most distinct from each other.

EEG connectivity (coherence) was calculated using built-in Fieldtrip [50] functions with the *mtmfft* method (which implements multitaper frequency transformation; taper: dpss) on 4-second segments, using the previously described bands of interest. Coherence values between two signals range from 0 to 1, with 1 indicating highly correlated signals. For statistical analysis, we selected a subset of electrodes from the standard 10–20 montage, commonly used in studies on AD patients [51,52]. However, the graphical results, including connectograms and matrix representations, are also displayed in the Supplementary Material for the high-density montage with 128 electrodes.

Functional MRI resting-state analysis was conducted using the CONN toolbox (version 21.a) [39]. Group-level ICA was used to identify 21 temporally coherent networks from the combined fMRI data of all subjects. The choice of 21 components is supported by prior research on individuals with various AD-related genetic polymorphisms [29], and using around 20 components is considered reasonable [25]. The selection of 21 components was further validated through visual inspection to ensure the distinctiveness of networks of interest. For subject-specific dimensionality reduction, a singular value decomposition of the z-score normalized BOLD signal was performed, with 64 components applied separately for each subject. Group-level analyses were conducted using a General Linear Model (GLM) [53]. The CONN software automatically assigned neural networks to components using the spatial match to template algorithm, which calculated the correlation between each group-level spatial map and CONN’s default networks with varying levels of spatial correlation coefficient. Although groups were matched by age and sex, these variables were included as nuisance regressors due to the significant age– and gender-related variability in rs-fMRI data [25]. Changes in the default mode network (DMN) are a characteristic feature of Alzheimer’s disease. Therefore, our primary focus was on this network, and we employed a hypothesis-driven approach to analyze the resting-state fMRI data, specifically examining the ICs with the highest correlation coefficient with the DMN.

### Statistics

Most statistical analyses were conducted using R [54] with RStudio [55] and custom scripts, with the exception of some neuroimaging statistics computed within specific toolboxes/software. Initially, we assessed the linearity between age and the dependent variables; since no linearity was assumed, age was not considered a covariate for statistics performed in R. For quantitative variables, either a two-way ANOVA test (with sex as a fixed factor) or a one-way ANOVA was used after verifying the assumptions of the tests. The normality of residuals was first assessed using the Shapiro-Wilk test and quantile (Q-Q) plots. Additionally, homoscedasticity was tested using Levene’s test (criterion: *p* < .05). For nominal variables, the chi-square test was employed. For ordinal variables and when ANOVA assumptions were not met, the non-parametric Kruskal-Wallis test was applied. ANOVAs were supplemented by standard post-hoc tests with Tukey correction for multiple comparisons. If the ANOVA test with Welch’s homogeneity correction was used, the Games-Howell post-hoc approach (with Tukey’s correction) was applied. For the Kruskal-Wallis test, Dunn’s post-hoc tests were used (with Holm correction). Statistical analysis of EEG coherence results was conducted in MATLAB using two-sample t-tests, with coherence calculated within the Fieldtrip toolbox. For t-test coherence results, FDR correction was applied using the Benjamini & Hochberg method [56] with a custom MATLAB function [57]. fMRI data statistics were derived from the SPM12 toolbox. Statistics on fMRI data connectivity were conducted at the cluster level, relying on parametric statistics derived from Gaussian Random Field theory [53,58]. The results were subjected to a thresholding approach involving a voxel-level threshold of *p* < .001 for cluster formation and a familywise corrected cluster-size threshold of *p*-FDR < .05 [59]. Statistical significance was defined as follows: a *p*-value ≤ .05 was considered significant, and a *p*-value > .05 and ≤ .09 was considered a trend. Where possible, all data were presented as mean (M) ± standard deviation (SD). Annotations on the figures are as follows: a *p*-value ≤ .05 is marked with *, < .01 with **, and < .001 with ***. Trend-level differences are marked with ∼ and the exact *p*-value.

## Results

### Characteristics of the participants

The study cohort comprised right-handed, middle-aged adults, with an equal distribution of females and males, ensuring gender balance in each subgroup (Tab. 1.). No significant differences between the groups regarding age, education, gender, handedness, possible alcohol problems, smoking status, health status were found (Tab. 1). Most respondents were generally healthy and were non-smokers. A+P– and A+P+ groups have more relatives (parents) with dementia than the N group (51.85% & 38.10% versus 25.81%), but there were no statistical differences.

The psychometric assessments indicated minor differences in depression/mood and self-esteem scales (BDI and SES, Tab. 1), with the A+P– group scoring worse than the N group (BDI: H(2) = 6.37, *p* < .05, post-hoc: A+P– vs. N *p* < .05; SES: F(2,76) = 3.17, *p* < .05, post-hoc: A+P– vs. N *p* < .05). Additionally, the A+P– group showed increased levels of neuroticism (NEO-NEU: H(2) = 6.04, *p* < .05, post-hoc: A+P– vs. N *p* < .05) and a tendency towards higher scores on the hopelessness scale in a stress coping strategies test (MINI-2 F(2,74) = 2.72, *p* = 0.07, post-hoc: A+P– vs. N *p* = .06). Given that a larger number of participants underwent psychometric testing and provided data on family history of dementia and demographics (N = 200 group), comparisons for significant findings in the neuroimaging phase groups were further analyzed in the this cohort, including the A-P+ group (not present in neuroimaging phase). To maintain consistent group structures based on allele assignment, individuals with the *APOE*-ε2 allele were excluded, leaving 174 participants. This resulted in unequal group sizes compared to the neuroimaging phase (A+P+ group, N = 21; A+P– group, N = 28; N group, N = 76; A-P+ group, N = 49). The extended comparison revealed significant differences in family history of dementia (X2(3) = 10.03, *p* < .05, N = 174, post–hoc: A+P– vs. N p < .05), with the A+P– group having more first-degree relatives with dementia than the N group. No significant differences were observed for BDI, SES, and the Mini-Cope-2 scale. Regarding neuroticism, differences were again identified (H(3) = 7.72, *p* = 0.05), but post-hoc tests considering six comparisons after introducing the A-P+ group revealed no significant differences after adjusting for multiple comparisons. Despite this, the A+P– group consistently had the highest neuroticism scores, while the N group had the lowest (A+P-: 20.29±9.42, A-P+: 20.41±8.77, A+P+: 15.62±8.18, N: 17.31±8.48).

### Power spectral measures

The A+P+ group demonstrated higher delta relative power than the N and/or A+P– groups both globally (F(2,72) = 4.19, *p* < .05, post-hoc: A+P– vs. A+P+ *p* < .05, A+P+ vs. N *p* < .05) and in certain electrode clusters (Fig. 2 and Tab. 2). There were no global differences in theta (*p* = .62), alpha-1 (*p* = .58), nor alpha-2 (*p* = .13) bands. The A+P+ group exhibited lower relative alpha-2 power in comparison to the N group in specific clusters (Fig. 2 and Tab. 2).

**Figure 2.**
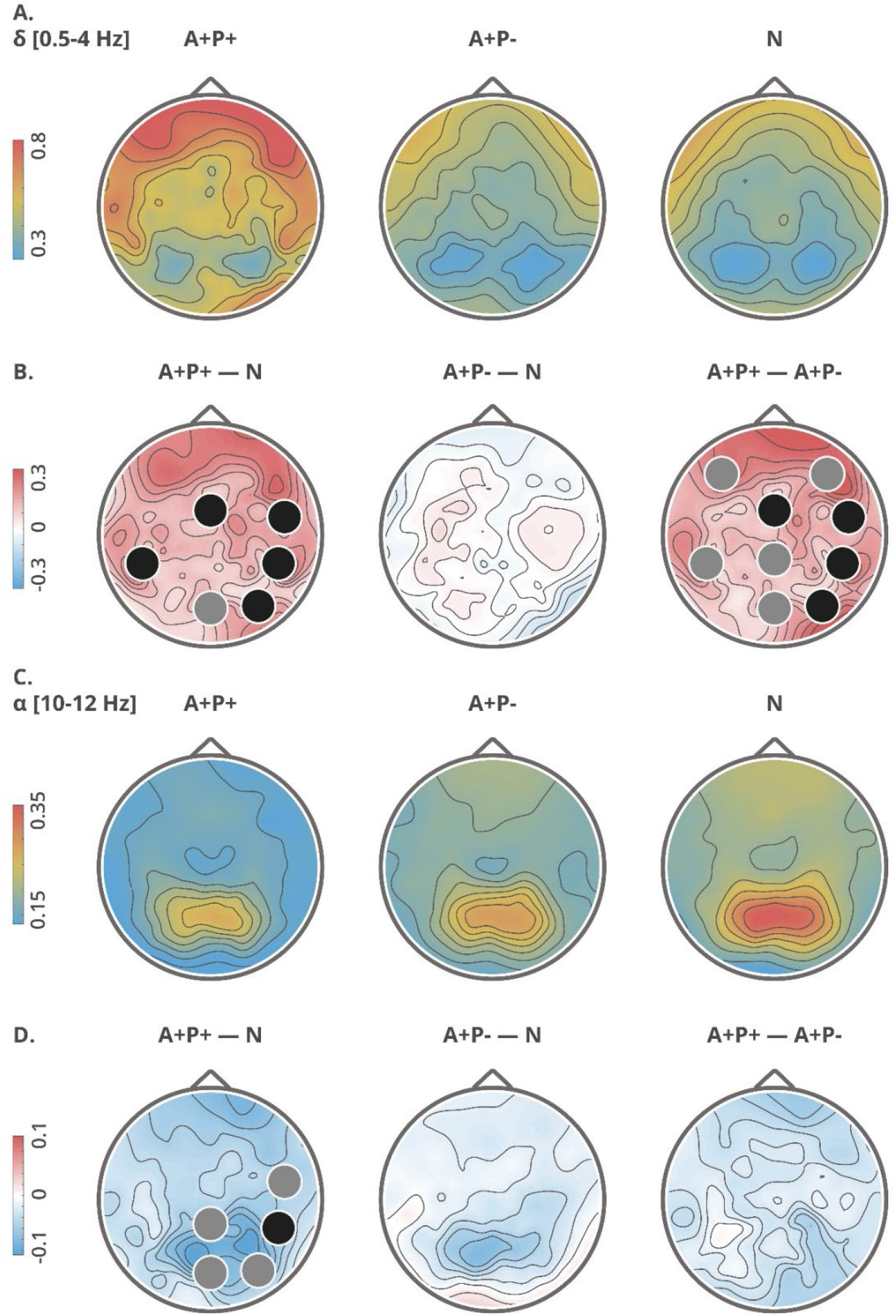
Topographical maps of delta (A, B) and alpha (C, D) relative power: group averages (upper rows: A, C) and group differences (lower rows: B, D). Clusters with significant differences (*p* < .05) between the groups are indicated with black circles, while trend-level differences (*p* < .09) are indicated with gray circles. The topographic maps are computed from high-density data, utilizing all 128 electrodes.

**Table 2.** Relative power for delta, theta, alpha-1 and alpha-2 bands: statistics, *p*-value and post-hoc details regarding group differences.

| Cluster | Statistic | $p$-value | Post-hoc details |
| --- | --- | --- | --- |
| **Delta** |  |  |  |
| MF | 1.21 | .31 | — |
| FL | 5.25 & | .07 | A+P- vs. A+P+ ($p = .085$) |
| FR | 5.80 & | .055 | A+P- vs. A+P+ ($p = .057$) |
| C | 3.70 | < .05 | A+P- vs. A+P+ ($p < .05$); A+P+ vs. N ($p < .05$) |
| CTL | 4.36 & | .11 | — |
| CTR | 4.79 | < .05 | A+P- vs. A+P+ ($p < .01$); A+P+ vs. N ($p < .05$) |
| PC | 2.51 | .09 | A+P- vs. A+P+ ($p = .09$) |
| PTL | 3.97 | < .05 | A+P- vs. A+P+ ($p = .053$);<br>A+P+ vs. N ($p < .05$) |
| PTR | 5.43 | < .01 | A+P- vs. A+P+ ($p < .01$); A+P+ vs. N ($p < .05$) |
| OC | 2.90 | .06 | A+P- vs. A+P+ ($p = .09$); A+P+ vs. N ($p = .09$) |
| OL | 3.11 & | .21 | — |
| OR | 4.80 | < .05 | A+P- vs. A+P+ ($p < .05$); A+P+ vs. N ($p < .05$) |
| **Theta** |  |  |  |
| MF | 0.21 | .81 | — |
| FL | 0.85 | .43 | — |
| FR | 0.58 | .57 | — |
| C | 0.06 | .95 | — |
| CTL | 0.72 | .49 | — |
| CTR | 0.65 | .53 | — |
| PC | 0.44 | .65 | — |
| PTL | 0.61 | .55 | — |
| PTR | 0.66 | .52 | — |
| OC | 0.63 | .54 | — |
| OL | 0.52 | .60 | — |
| OR | 0.54 | .58 | — |
| **Alpha-1** |  |  |  |
| MF | 0.51 # | .60 | — |
| FL | 0.69 | .51 | — |
| FR | 1.34 | .27 | — |
| C | 0.39 # | .68 | — |
| CTL | 0.35 | .71 | — |
| CTR | 0.81 | .45 | — |
| PC | 0.53 | .59 | — |
| PTL | 0.35 | .70 | — |
| PTR | 0.47 | .63 | — |
| OC | 0.65 | .53 | — |
| OL | 0.61 | .55 | — |
| OR | 0.49 | .62 | — |
| **Alpha-2** |  |  |  |
| MF | 1.40 # | .25 | — |
| FL | 2.04 & | .36 | — |
| FR | 3.35 & | .19 | — |
| C | 1.64 | .20 | — |
| CTL | 1.12 | .33 | — |
| CTR | 3.09 | .052 | A+P+ vs. N ($p < .05$) |
| PC | 2.70 | .07 | A+P+ vs. N ($p = .08$) |
| PTL | 1.99 | .15 | — |
| PTR | 3.27 # | <.05 | A+P+ vs. N ($p < .05$) |
| OC | 3.02 | .055 | A+P+ vs. N ($p = .053$) |
| OL | 2.15 | .12 | — |
| OR | 2.70 | .07 | A+P+ vs. N ($p = .06$) |

NOTE. $p$ -value: group factor. Post-hoc details are provided only for significant or trend level differences. #, one-way ANOVA; &, Kruskal-Wallis test; MF, midfrontal; FL, frontal-left; FR, frontal-right; C, central; CTL, central-left; CTR, central-right; PC, posterior-central; PTL, posterior-left; PTR, posterior-right; OC, occipital-central; OL, occipital-left; OR, occipital-right.

Male participants were consistently characterized by higher theta relative power than females, both globally (F(1,72) = 5.90, *p* < .05) and in certain electrode clusters (Supplementary Table 2). They were also characterized by lower high alpha band power than females in certain clusters (Supplementary Table 2). Additionally, there was an interaction between sex and group regarding delta relative power (Supplementary Figure 1), indicating that males in the A+P+ group had higher delta power than males or females in the N or A+P–groups, both globally (F(2,72) = 3.17, *p* < .05, post-hoc: M A+P-vs. M A+P+ *p* < .01, M A+P+ vs. M N *p* < .05, M A+P+ vs. F N *p* < .05, F A+P-vs. M A+P+ *p* = .08) and in specific electrode clusters (Supplementary Table 1). A statistically significant interaction was also observed between the group and sex factors on lower alpha relative power, both globally (F(2,72) = 5.84, *p* < .01, post-hoc: F A+P-vs. M A+P-*p* < .05, M A+P-vs. M N *p* < .05, M A+P-vs. F A+P+ *p* = .07) and in specific electrode clusters (Supplementary Figure 1). Female participants from the A+P+ group had lower low alpha relative power than males in the A+P-group, and males from the N group had lower low alpha relative power compared to males in the A+P-group. Additionally, females in the A+P-group exhibited lower low alpha relative power than males in the same group.

### Signal complexity

The signal complexity results are depicted in Figure 3, showing the average global HFD plotted against the tuning parameter *k*_max_. The A+P+ group demonstrated significantly lower signal complexity (M ± SD, 1.83 ± 0.04) compared to the N group (M ± SD, 1.85 ± 0.03), and showed a trend towards lower complexity than the A+P-group (M ± SD, 1.85 ± 0.03) globally (F(2,72) = 5.23, *p* < 0.01, post-hoc: A+P+ vs. N *p* < .01, A+P+ vs. A+P-*p* = .07) and in specific clusters (Tab. 3 and Fig. 3). Global average HFD in relation to *k*_max_ is shown in Supplementary Figure 2.

**Figure 3.**
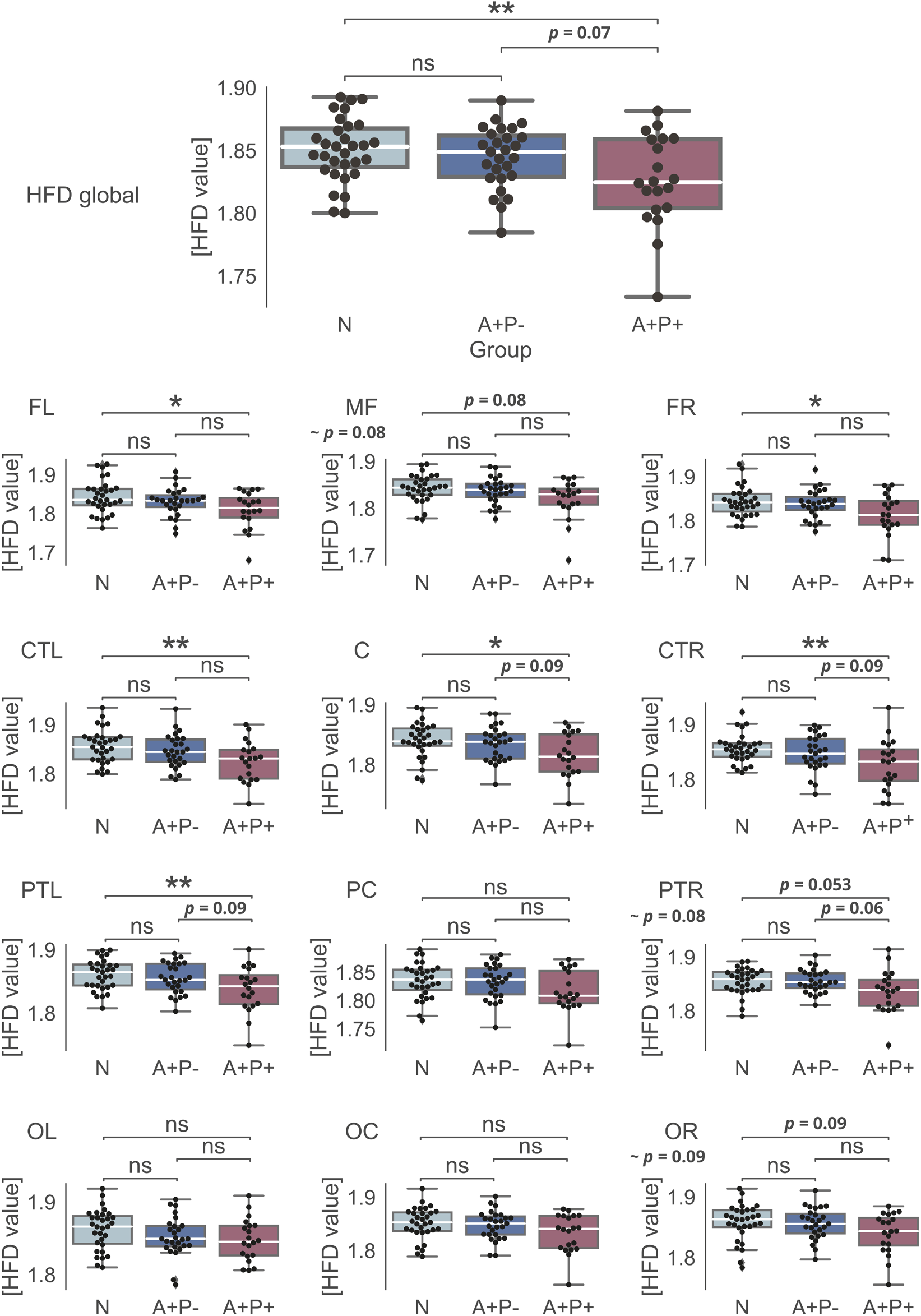
Average Higuchi’s Fractal Dimension. Differences between the groups on *k*_max_ = 82.

**Table 3.** HFD values. Descriptive statistics and group differences.

| Clusters | Group ($M \pm SD$ values) | | | $p$-value | Statistic | Post-hoc details |
| --- | --- | --- | --- | --- | --- | --- |
|  | A+P+ | A+P- | N |  |  |  |
| MF | $1.82 \pm 0.04$ | $1.84 \pm 0.03$ | $1.84 \pm 0.03$ | .08 | 5.00 & | A+P+ vs. N ($p < .08$) |
| FL | $1.81 \pm 0.05$ | $1.83 \pm 0.04$ | $1.84 \pm 0.04$ | $< .05$ | 4.36 | A+P+ vs. N ($p < .05$) |
| FR | $1.81 \pm 0.05$ | $1.84 \pm 0.03$ | $1.84 \pm 0.03$ | $< .05$ | 4.70 | A+P+ vs. N ($p < .05$) |
| C | $1.82 \pm 0.04$ | $1.84 \pm 0.03$ | $1.83 \pm 0.03$ | $< .05$ | 4.44 | A+P+ vs. N ($p < .05$);<br>A+P+ vs. A+P- ($p = .09$) |
| CTL | $1.83 \pm 0.04$ | $1.85 \pm 0.03$ | $1.86 \pm 0.03$ | $< .05$ | 4.69 | A+P+ vs. N ($p < .01$) |
| CTR | $1.83 \pm 0.05$ | $1.85 \pm 0.03$ | $1.86 \pm 0.03$ | $< .01$ | 5.13 | A+P+ vs. N ($p < .01$);<br>A+P+ vs. A+P- ($p < .09$) |
| PC | $1.82 \pm 0.04$ | $1.83 \pm 0.03$ | $1.84 \pm 0.03$ | .10 | — | — |
| PTL | $1.84 \pm 0.04$ | $1.86 \pm 0.02$ | $1.86 \pm 0.02$ | $< .05$ | 4.70 | A+P+ vs. N ($p < .01$);<br>A+P+ vs. A+P- ($p < .09$) |
| PTR | $1.84 \pm 0.04$ | $1.86 \pm 0.02$ | $1.86 \pm 0.02$ | .08 | 3.43 # | A+P+ vs. N ($p = .053$);<br>A+P+ vs. A+P- ($p = .06$) |
| OC | $1.83 \pm 0.04$ | $1.85 \pm 0.03$ | $1.85 \pm 0.03$ | .10 | — | — |
| OL | $1.85 \pm 0.03$ | $1.85 \pm 0.03$ | $1.86 \pm 0.03$ | .13 | — | — |
| OR | $1.84 \pm 0.03$ | $1.86 \pm 0.03$ | $1.86 \pm 0.03$ | .09 | 2.5 # | A+P+ vs. N ($p = .09$) |

NOTE. The bolded font indicate a significant or trend result. $p$ -value is shown for *Group* factor. #, One-way ANOVA; &, Kruskal-Wallis test; MF, midfrontal; FL, frontal-left; FR, frontal-right; C, central; CTL, central-left; CTR, central-right; PC, posterior-central; PTL, posterior-left; PTR, posterior-right; OC, occipital-central; OL, occipital-left; OR, occipital-right.

Moreover, sex had a significant impact on HFD values, with males displaying reduced signal complexity both globally (F(1,72) = 4.08, *p* < .05) and in certain electrode clusters (Supplementary Table 2) (CTR cluster: F(1,72) = 5.46, *p* < .05; PTL: F(1,72) = 6.81, *p* < .05; and a trend level in the C cluster: F(1,72) = 3.69, *p* = 0.06). An interaction between sex and group factors was found in clusters summarized in Supplementary Table 4: FL and FR and at a trend level (p = .06) in the CTR cluster and globally. These findings mirrored the main results for the group factor alone, with females/males from the A+P+ group exhibiting the lowest HFD values compared to females/males from the A+P– and N groups (which had the highest values).

### Connectivity

The analysis of EEG connectivity, measured by the coherence of frequency components (bands), identified subtle patterns indicating some differences between the groups at the *p*-uncorrected level. In the delta band, the A+P+ group displayed greater coherence compared to the A+P– and N groups. Furthermore, the A+P-group showed reduced theta coherence relative to the N group. Both the A+P+ (low and high alpha) and A+P– (low alpha) groups exhibited (mostly) decreased coherence in the alpha band compared to the N group (as depicted in Supplementary Figure 3, with uncorrected *p*-values below the alpha .05 level). However, it is crucial to highlight that none of these associations remained significant after applying the false discovery rate (FDR) correction. Additionally, Supplementary Figure 4 shows coherence results using a high-density montage with all 128 electrodes, including connectograms and difference matrix representations for a thorough depiction of global connectivity.

The analysis of fMRI ICA-based connectivity revealed some subtle differences between the studied groups. The correlation of 21 estimated components with known networks is detailed in Supplementary Table 5, showing the best three matching components: IC 11 (r = .38), IC 5 (r = .34), and IC 20 (r = 0.11). Their representations are shown in Supplementary Figure 5. While IC 5 and IC 11 clearly depict posterior parts of the DMN, IC 20 shows mixed areas, which is explained by only an r = .11 correlation coefficient with the DMN. Additionally, IC 20 includes signals from the cerebrospinal fluid (CSF) and ventricular regions (cyan color). As indicated by the correlation coefficients, these components do not perfectly align with traditional DMN areas and also include other, nearby regions. Two DMN related components (IC 11 and IC 5) were selected for further group comparisons.

The A+P-group exhibited significantly lower network strength in a cluster in the right temporo-occipital part of the middle temporal gyrus (toMTG, IC11) compared to the N group (Tab 4. And Fig. 4). This effect was significant at *p*-FDR < .01. No significant differences were found in IC 5 between the groups.

**Table 4.** Spatial cluster (location, size, and anatomical alignment) associated with IC 11, highlighting significant difference between the groups.

| Group comparison | IC | Cluster (x, y, z) | $k$ | $p$-FDR < | Label | % of voxels in the cluster covering % of labeled area |
| --- | --- | --- | --- | --- | --- | --- |
| A+P- – N | 11 | 62, -44, 6 | 141 | .01 | toMTG r | 98% / 12% |

NOTE. r, right; $k$ , cluster size; toMTG, middle temporal gyrus temporo-occipital part.

**Figure 4.**
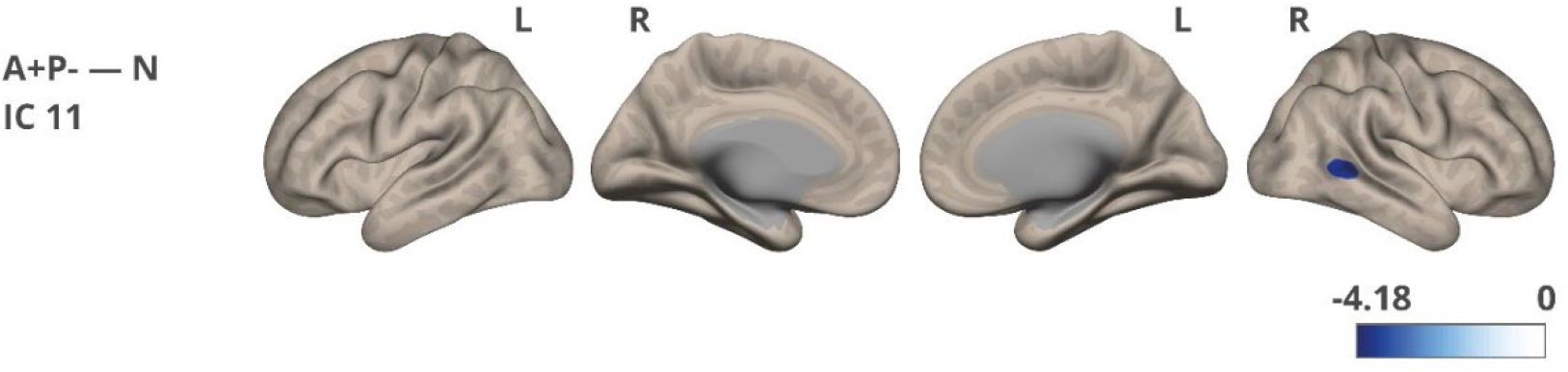
Network strength/connectivity differences were noted between the A+P– and N groups for IC 11. The color corresponds to the direction of the difference and obtained t-value. The images were visualized using the CONN toolbox, with t-values adjusted to fit the cortical structure on a glass ICBM template brain surface display, presented in lateral and medial views.

## Discussion

We examined EEG and fMRI resting-state data from healthy middle-aged individuals with various alleles of the Alzheimer’s disease risk genes *APOE* and *PICALM*. The groups were demographically balanced, and we discovered that risk carriers exhibit some AD hallmarks in domains directly (EEG, fMRI) and indirectly (psychometric evaluation) associated with brain functioning.

The entire cohort consisted of well-functioning individuals from major urban areas who were professionally active and had a high level of education. The groups performed equally well on both the intelligence test (RAVEN) and the California Verbal Learning Test (CVLT) which assesses verbal learning and memory. Previously reported CVLT results for healthy but much older *APOE* carriers showed that they either did not differ from no-risk groups [60] or exhibited a higher frequency of recall intrusion errors [61]. Despite the lack of clear cognitive impairment, the subjects displayed group differences in specific psychological characteristics. Participants with a single risk factor (A+P-) declared worse psychological well-being compared to the no-risk group. They exhibited lower self-esteem, higher scores on the BDI scale (related to depression and lowered mood), heightened levels of neuroticism, and a tendency to use less effective stress coping strategies. It is well established that factors such as depression, lowered mood, and neuroticism act as risk factors for Alzheimer’s disease and may interact with *APOE* in provoking the disease [2,61–64]. The BDI scores differentiated the groups, although the average scores were within the normal range. Prolonged stress, which can cause these psychological symptoms, is also described as a risk factor [63]. Specifically, midlife stress is associated with the development of dementia in later life [65]. Notably, these differences were significant primarily for the cohort that underwent neuroimaging testing. However, for the entire cohort of 200 individuals recruited for genetic screening, statistical significance was achieved only for neuroticism.

In double risk carriers (A+P+ group), EEG resting state was characterized by two of the most noticeable and sensitive markers of AD: a shift of the power spectrum to lower frequencies (known as “slowing of the EEG”) and decreased signal complexity. EEG “slowing” was evidenced by higher delta and lower alpha relative power, while a lower Higuchi fractal dimension indicated reduced signal complexity. These two measures, complexity and EEG dynamics, are strongly related [66] and correspond to the capacity for information processing (i.e., less complex signal = lower capacity), which changes due to neurodegeneration as well as during natural stages of brain development, maturation, and aging. Additionally, the impact of common genetic variations (e.g., *APOE* gene) on cognitive/brain functioning increases with aging [67]. EEG abnormalities in normal aging also include changes in spectral content, such as decreased power in delta, theta, and alpha peak frequencies [13] and decline in complexity (significantly affecting central-parietal areas, especially right-shifted clusters) [47]. In MCI patients, research has shown a small increase in the power of delta and theta bands in temporal areas [13]. Upper alpha, but not lower alpha power, was distinguishable among controls and AD patients in another study [15]. AD patients with *APOE* risk have stronger EEG “slowing” than carriers of neutral *APOE* alleles [4,68], although one study showed opposite results [69]. Power perturbations within the temporal and parietal areas are especially sensitive indicators for distinguishing Alzheimer’s disease patients from healthy controls [18]. A reduction in complexity in AD patients is clearly noticeable, particularly affecting the temporal-occipital regions [47]. The EEG “slowing” marker is believed to be associated with neuronal loss, axonal pathology, and cholinergic deficits, which affect functional connections in the cortex [20]. Decreased signal complexity is also related to either neuronal loss or neurotransmitter deficiencies, such as acetylcholine [20]. Atrophy in cholinergic neurons may be the primary source of EEG “slowing,” as these neurons are most affected by AD. The cholinergic hypothesis of AD and memory dysfunction in the elderly was proposed over 50 years ago [20,70,71]. The cholinergic system regulates various aspects of brain function—cognition, locomotion, attention, sleep, arousal, and sensory processing—by modulating neuronal activity via acetylcholine receptors. Cholinergic drugs tend to reverse EEG “slowing,” supporting this hypothesis [20,72]. Anticholinergic drugs (e.g., scopolamine), which block the stimulation of post-synaptic receptors, cause EEG “slowing” [20,72,73]. *APOE*-ε4 positive AD patients are characterized by more severe cholinergic deficits than patients with a neutral *APOE* genotype [74].

In our study, the groups did not exhibit significant differences in connectivity, as measured by EEG coherence at the FDR-corrected level. However, on the uncorrected level, they showed subtle trends in the same direction as reported among AD patients: decrease in alpha coherence in the at-risk (A+P+ or A+P-) groups compared to the N group, and increased delta coherence in the at-risk A+P+ group compared to the A+P– and N groups. A recent systematic review showed that in 24 out of 34 studies comparing AD patients to healthy controls, AD patients had significantly decreased coherence within the alpha band [75]. The results for coherence in lower frequencies (< 7 Hz) were less consistent and less frequently significant [75]. Generally, alpha coherence tends to be decreased in AD patients [13], while delta and theta coherences tend to be increased compared to matching controls [13].

Using an ICA-based approach, we identified small yet significant differences in fMRI connectivity. This analysis primarily focused on independent components linked to the DMN, which is often impaired in Alzheimer’s disease patients [25,76]. The A+P-group (compared to the N group) exhibited significantly reduced network strength in a cluster encompassing the right temporo-occipital part of the middle temporal gyrus. Although the MTG is not traditionally considered a core component of the DMN, numerous studies demonstrate its involvement within the posterior DMN and discuss DMN role in semantic cognition [77–79]. Additionally, the medial temporal lobe is recognized as one of the first regions affected by Alzheimer’s disease, showing early signs of atrophy and the presence of neurofibrillary tangles [80,81]. Network changes in *APOE* risk-carries (without amyloid burden) were previously reported in the literature [31,82]. Certain studies have revealed decreased connectivity within the posterior DMN in older *APOE*-ε4 carriers (70-89 years old) compared to non-carriers [83], while no effect was observed in middle-aged adults [32]. Research has consistently indicated decreased connectivity within the posterior DMN in individuals with MCI and Alzheimer’s disease patients, while recently showing increased connectivity within the frontal parts of the DMN [25,76]. The causes of network changes are still unclear; it is unknown whether they are related to amyloid deposition, whether they represent a compensatory mechanism in response to amyloid atrophy and toxicity, or to what extent they are influenced by genetic factors [25]. Atrophy and hypometabolism are known to be partially responsible for observable network changes in amyloid-positive individuals [84]. The concept of network-based functional compensation suggests that alterations in the brain’s functional architecture, influenced by genes such as *APOE* and *PICALM*, may enable the brain to adapt and compensate for changes or disruptions in specific brain regions or networks.

In summary, our findings reinforce previous research and suggest that *APOE* and *PICALM* shape the functional architecture of the resting brain even in the absence of dementia. Future research should prioritize studying non-demented risk carriers throughout their lifespan to understand the impact of these genetic variations on aging and to uncover the biological mechanisms underlying their association with neurodegenerative diseases. Early detection of Alzheimer’s should employ multimodal approaches that consider genetic burden (such as the *APOE* and *PICALM* risks in our study), and additionally some or all markers like: blood-based biomarkers, MRI/fMRI/EEG abnormalities, cognitive performance, health, lifestyle, demographic factors, and neuropsychological assessments. Together, these markers may help identify individuals at risk of developing dementia in the future, allowing for potential early and successful interventions.

## Authors’ Contributions

PD designed and implemented the experiment, collected the data, performed data analysis and statistical analysis, prepared figures, and wrote the original draft of the manuscript. JW carried out the first and second stages of the MRI/fMRI data analysis and revised the final manuscript draft. TW supervised the fMRI data analysis and revised the final manuscript draft. EK designed the experiment, reviewed, and revised the final version of the manuscript. All authors read and approved the final manuscript version.

## Acknowledgments

We thank participants for their valuable time and willingness to participate in the study. We appreciate help of Ingrida Antonova and Olga Stefańska in EEG acquisition.

## Supplementary Material

**Supplementary Figure 1.**
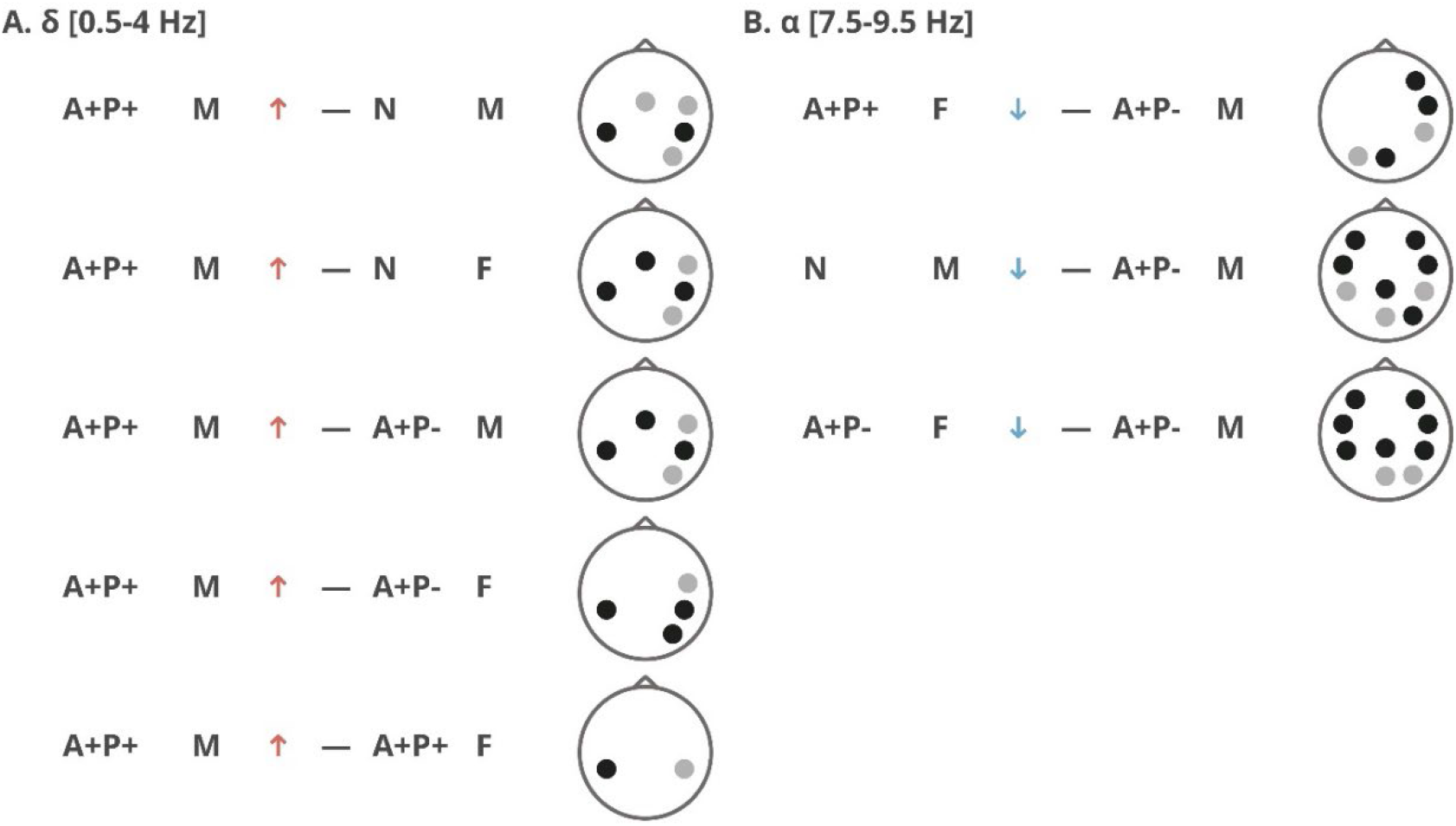
A graphical summary illustrating the interaction between group and sex factors in the EEG power spectrum results: delta relative power (A) and low alpha relative power (B). An upward arrow signifies higher power for the first reported group, and a downward arrow signifies lower power. The topographic maps indicate the locations of clusters with significant and near-significant effects. Clusters with significant differences (p < 0.05) are marked with black circles, while trend-level differences (p < 0.09) are marked with gray circles

**Supplementary Table 1.**
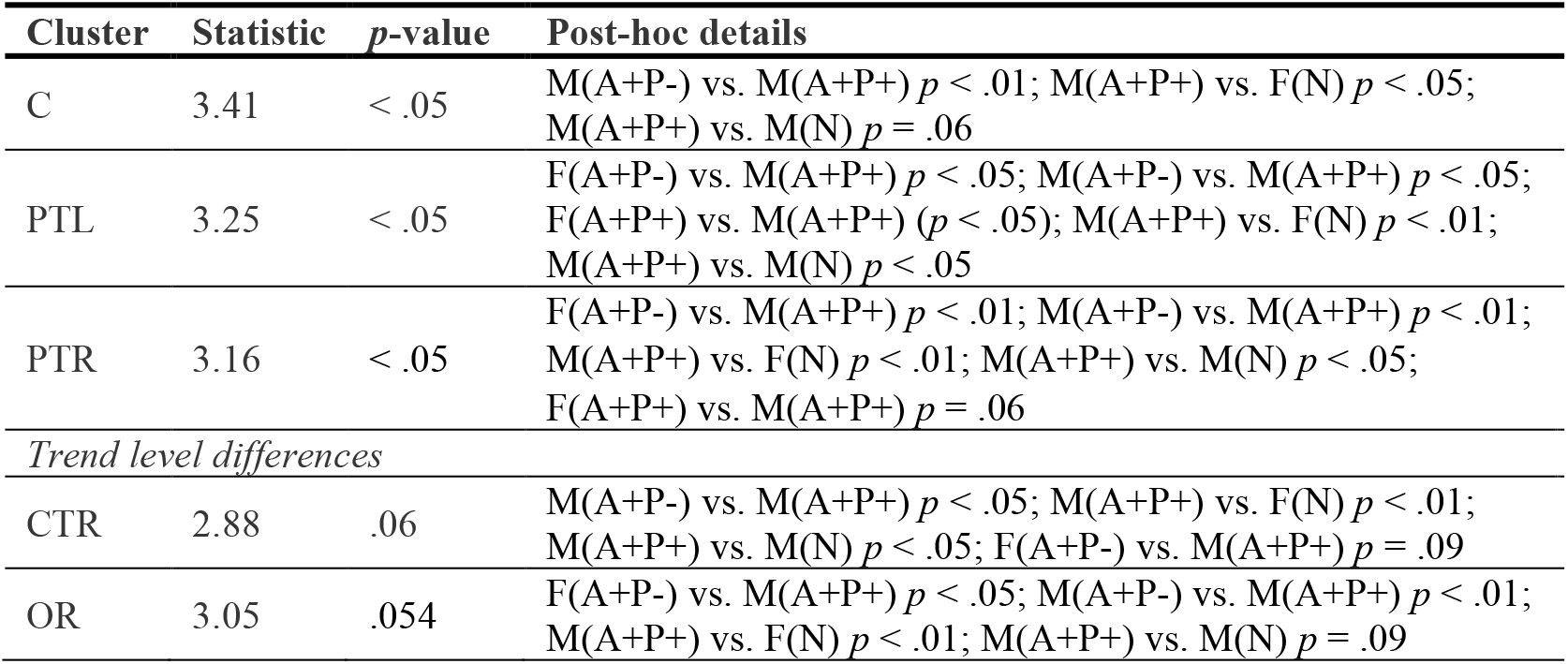
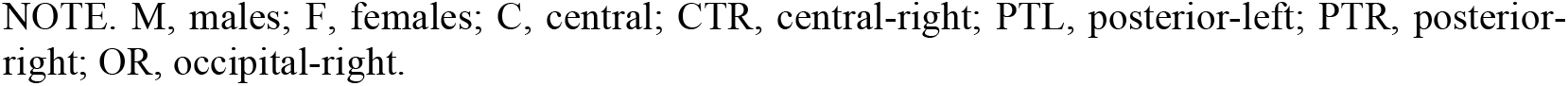
Relative power within the delta band: statistical values and post-hoc information related to sex*group interaction differences that are either significant or indicate a trend.

| Cluster | Statistic | *p*-value | Post-hoc details |
| --- | --- | --- | --- |
| C | 3.41 | < .05 | M(A+P-) vs. M(A+P+) $p < .01$; M(A+P+) vs. F(N) $p < .05$; M(A+P+) vs. M(N) $p = .06$ |
| PTL | 3.25 | < .05 | F(A+P-) vs. M(A+P+) $p < .05$; M(A+P-) vs. M(A+P+) $p < .05$; F(A+P+) vs. M(A+P+) ($p < .05$); M(A+P+) vs. F(N) $p < .01$; M(A+P+) vs. M(N) $p < .05$ |
| PTR | 3.16 | < .05 | F(A+P-) vs. M(A+P+) $p < .01$; M(A+P-) vs. M(A+P+) $p < .01$; M(A+P+) vs. F(N) $p < .01$; M(A+P+) vs. M(N) $p < .05$; F(A+P+) vs. M(A+P+) $p = .06$ |
| *Trend level differences* |  |  |  |
| CTR | 2.88 | .06 | M(A+P-) vs. M(A+P+) $p < .05$; M(A+P+) vs. F(N) $p < .01$; M(A+P+) vs. M(N) $p < .05$; F(A+P-) vs. M(A+P+) $p = .09$ |
| OR | 3.05 | .054 | F(A+P-) vs. M(A+P+) $p < .05$; M(A+P-) vs. M(A+P+) $p < .01$; M(A+P+) vs. F(N) $p < .01$; M(A+P+) vs. M(N) $p = .09$ |

NOTE. M, males; F, females; C, central; CTR, central-right; PTL, posterior-left; PTR, posterior-right; OR, occipital-right.

**Supplementary Table 2.** Relative power within the theta and higher alpha band and HFD: statistical values and post-hoc information related to the sex factor.

| Cluster | Theta<br>(Statistic) | Theta<br>(*p*-value) | Alpha-2<br>(Statistic) | Alpha-2<br>(*p*-value) | HFD<br>(Statistic) | HFD<br>(*p*-value) |
| --- | --- | --- | --- | --- | --- | --- |
| MF | 4.01 | < .05 | — | — | — | — |
| FL | 4.73 | < .05 | — | — | 0.02 | .88 |
| FR | 4.04 | < .05 | — | — | — | .21 |
| C | 5.21 | < .05 | 3.19 | .08 | 3.69 | .06 |
| CTL | 4.13 | < .05 | 4.89 | < .05 | 0.87 | .36 |
| CTR | 6.10 | < .05 | 4.00 | < .05 | 5.46 | < .05 |
| PC | 3.56 | .06 | 1.53 | .22 | 2.65 | .11 |
| PTL | 5.90 | < .05 | 4.47 | < .05 | 6.81 | < .05 |
| PTR | 5.65 | < .05 | — | — | — | — |
| OC | 4.99 | < .05 | 6.10 | < .05 | 2.87 | .10 |
| OL | 10.48 | < .01 | 8.45 | < .01 | 8.03 | < .01 |
| OR | 8.10 | < .01 | 8.48 | < .01 | — | — |

NOTE. MF, midfrontal; FL, frontal-left; FR, frontal-right; C, central; CTL, central-left; CTR, central-right; PC, posterior-central; PTL, posterior-left; PTR, posterior-right; OC, occipital-central; OL, occipital-left; OR, occipital-right.

**Supplementary Table 3.** Relative power within the lower alpha band: statistical values and post-hoc information related to sex*group interaction differences that are either significant or indicate a trend.

| Cluster | Statistic | *p*-value | Post-hoc details |
| --- | --- | --- | --- |
| FL | 5.76 | < .01 | F(A+P-) vs. M(A+P-) $p < .05$; M(A+P-) vs. M(N) $p < .05$ |
| FR | 5.65 | < .01 | F(A+P-) vs. M(A+P-) $p < .05$; M(A+P-) vs. F(A+P+) $p < .05$; M(A+P-) vs. M(N) ($p < .05$) |
| CTL | 5.75 | < .01 | F(A+P-) vs. M(A+P-) $p < .05$; M(A+P-) vs. M(N) $p < .05$ |
| CTR | 5.44 | < .01 | F(A+P-) vs. M(A+P-) $p < .05$; M(A+P-) vs. F(A+P+) $p < .05$; M(A+P-) vs. M(N) $p < .05$ |
| PC | 5.93 | < .01 | F(A+P-) vs. M(A+P-) $p < .05$; M(A+P-) vs. M(N) $p < .05$ |
| PTL | 5.96 | < .01 | F(A+P-) vs. M(A+P-) $p < .05$; M(A+P-) vs. M(N) $p = .07$ |
| PTR | 5.00 | < .01 | F(A+P-) vs. M(A+P-) $p < .05$; M(A+P-) vs. F(A+P+) $p = .09$; M(A+P-) vs. M(N) $p = .08$ |
| OC | 5.29 | < .01 | F(A+P-) vs. M(A+P-) $p = .07$; M(A+P-) vs. F(A+P+) $p < .05$; M(A+P-) vs. M(N) $p = .08$ |
| OL | 5.11 | < .01 | M(A+P-) vs. F(A+P+) $p = .07$ |
| OR | 5.76 | < .01 | F(A+P-) vs. M(A+P-) $p = .07$; M(A+P-) vs. M(N) $p < .05$ |

NOTE. M, males; F, females; FL, frontal-left; FR, frontal-right; CTL, central-left; CTR, central-right; PC, posterior-central; PTL, posterior-left; PTR, posterior-right; OC, occipital-central; OL, occipital-left; OR, occipital-right.

**Supplementary Table 4.** Statistical values and post-hoc information related to sex*group interaction in HFD analysis, highlighting significant or trend-level results.

| Cluster | p-value | Statistic | Post-hoc details |
| --- | --- | --- | --- |
| FL | < .01 | 5.41 | M(N) > M(A+P+) $p < .001$; M(A+P+) < M(A+P-) $p < .05$; F(A+P+) > M(A+P+) $p = .06$; F(N) > M(A+P+) $p < .05$ |
| FR | < .01 | 5.03 | F(N) > M(A+P+) $p < .05$; M(N) > M(A+P+) $p < .001$; F(A+P+) > M(A+P+) $p < .05$; M(A+P+) < F(A+P-) $p < .01$; M(A+P+) < M(A+P-) $p < .05$ |
| *Trend level differences* |  |  |  |
| Global | .06 | 2.92 | F(N) > M(A+P+) $p < .01$; M(N) > M(A+P+) $p < .01$; F(A+P+) > M(A+P+) $p < .05$; M(A+P+) < F(A+P-) ($p < .01$) |
| CTR | .06 | 2.97 | F(N) > M(A+P+) $p < .01$; M(N) > M(A+P+) $p < .01$; F(A+P+) > M(A+P+) ($p < .05$); M(A+P+) < F(A+P-) ($p < .01$); M(A+P+) < M(A+P-) $p = .07$ |

NOTE. M, males; F, females; FL, frontal-left; FR, frontal-right; CTR, central-right.

**Supplementary Figure 2.**
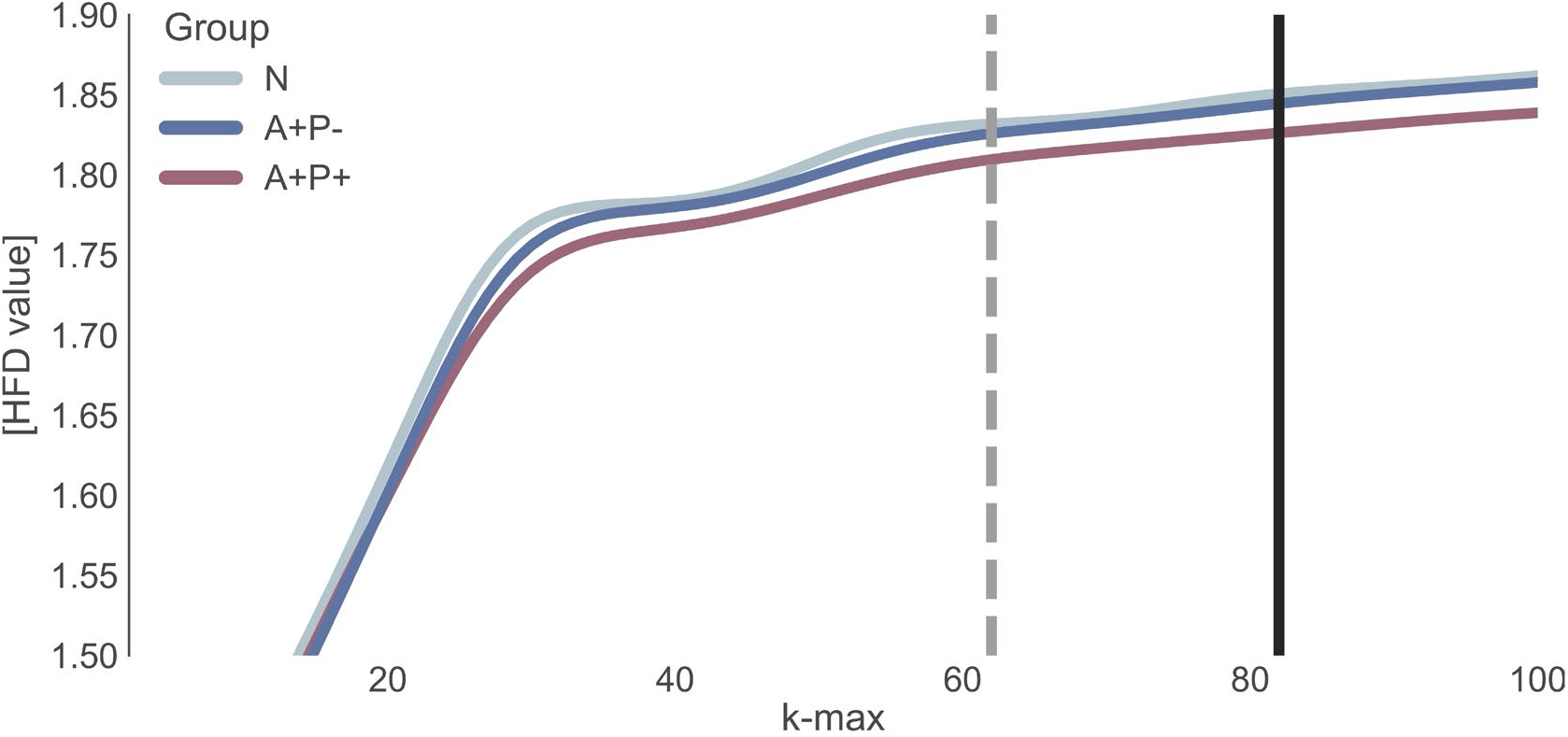
Dependence of Higuchi’s Fractal Dimension value on the parameter *k*_max_. The dotted line indicates the start of HFD stability, and the black line indicates the *k*_max_ selected for the between-group comparison of the HFD measure

**Supplementary Figure 3.**
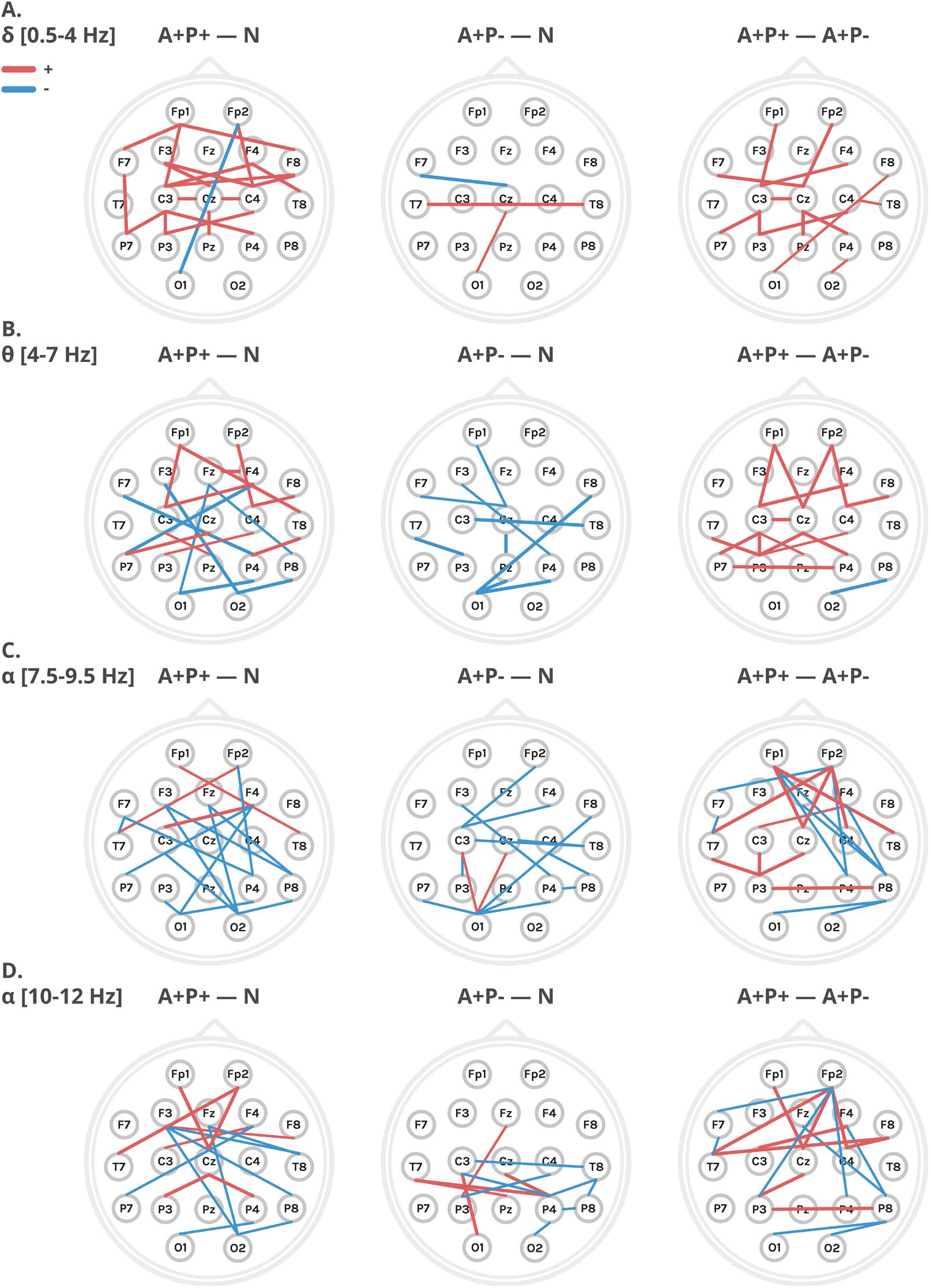
Differences in connectivity, measured by coherence, were analyzed across the 19 electrodes in the classical 10-20 montage for each frequency band. Red lines represent higher coherence for the first group compared to the second group (e.g., A+P+ versus N), while blue lines represent lower coherence. The image displays only significant t-test results (Fieldtip) between the electrodes, but none of these retained significance after FDR correction.

**Supplementary Figure 4.**
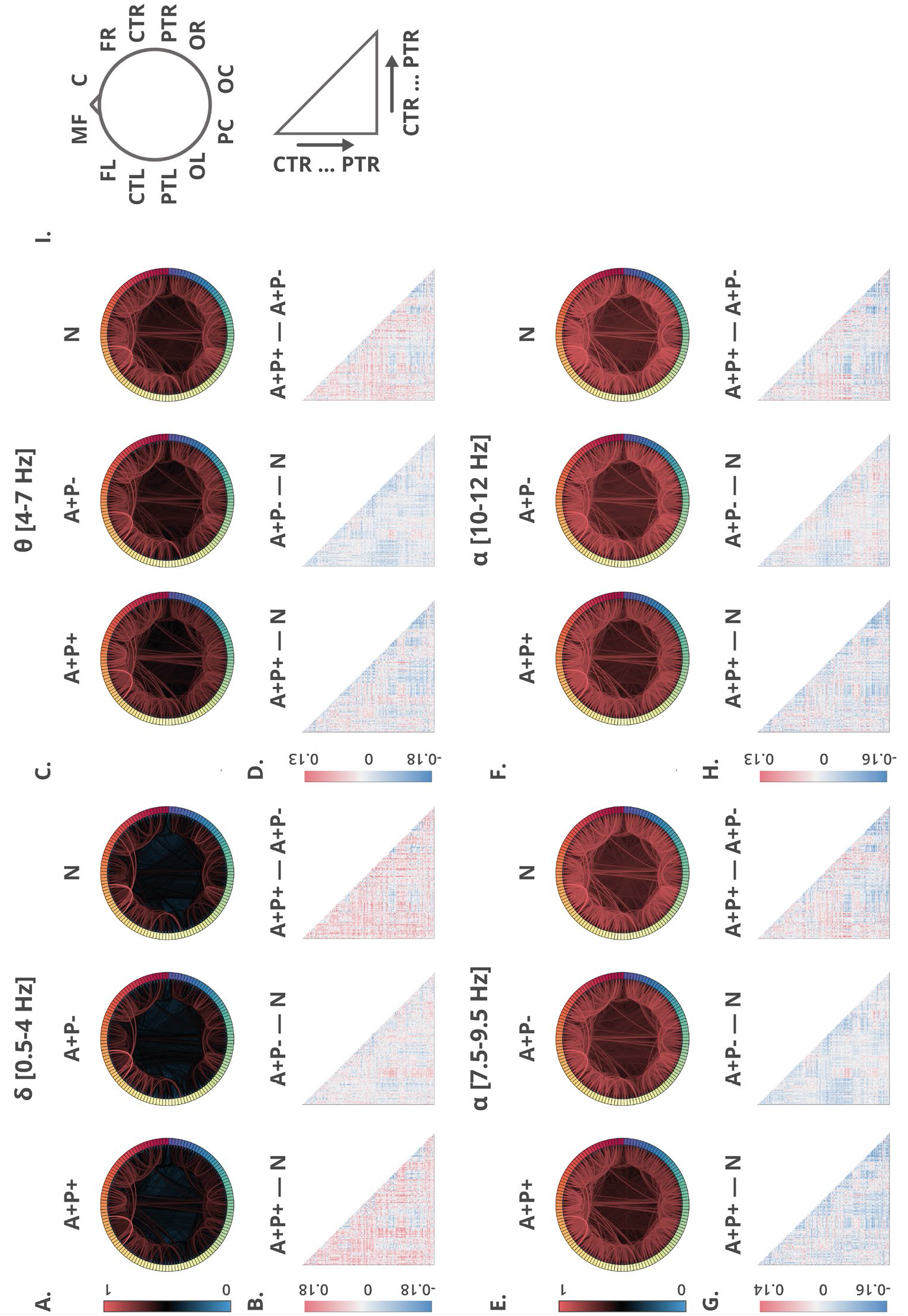
EEG connectivity (coherence) using a high-density montage with all 128 electrodes. Connectograms for each group are shown, with distinct frequency bands in panels labeled as A, E, C, and F in the graphs. Matrix representations highlighting group differences are displayed in graphs B, D, G, and H. Graph I serves as a legend, detailing electrode placement in all plots, arranged according to previously defined clusters.

**Supplementary Table 5.** Mapping of the 21 Independent Components to neural networks.

| Neural network | Independent component No and its correlation coefficient with the given network |
| --- | --- |
| Default Mode Network | **11 ($r=0.38$)**, 5 ($r=0.34$), 20 ($r=0.11$) |
| Sensorimotor Network | **21 ($r=0.46$)**, 16 ($r=0.43$), 19 ($r=0.24$) |
| Visual Network | **12 ($r=0.56$)**, 15 ($r=0.41$), 13 ($r=0.25$) |
| Saliency Network | **2 ($r=0.35$)**, 10 ($r=0.10$), 14 ($r=0.08$) |
| Dorsal Attention Network | **19 ($r=0.38$)**, 10 ($r=0.27$), 13 ($r=0.10$) |
| Fronto-Parietal Network | **6 ($r=0.20$)**, 17 ($r=0.17$), 4 ($r=0.14$) |
| Language Network | **3 ($r=0.39$)**, 8 ($r=0.21$), 10 ($r=0.11$) |
| Cerebellar Network | **1 ($r=0.37$)**, 18 ($r=0.34$), 17 ($r=0.20$) |

**Supplementary Figure 5.**
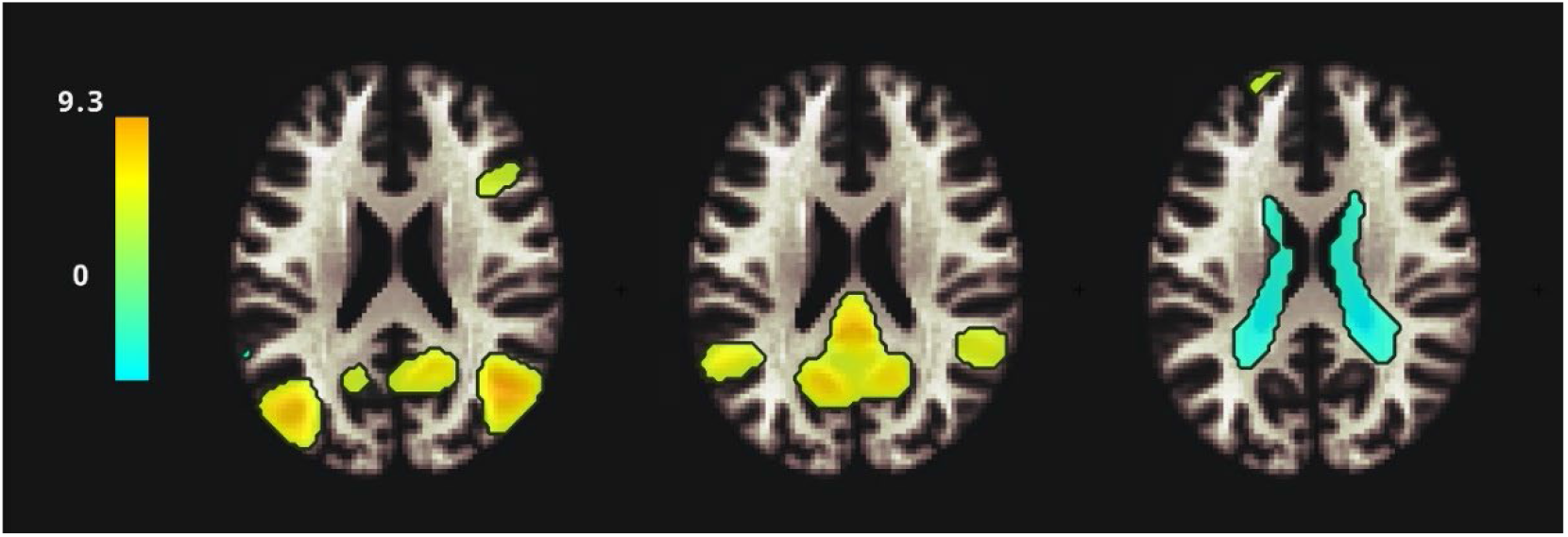
The representation of IC 5, IC 11, and IC 20 – components with the highest correlation coefficients with the Default Mode Network (DMN). Retrieved from the CONN toolbox.

